# Potential anti-monkeypox virus activity of atovaquone, mefloquine, and molnupiravir, and their potential use as treatments

**DOI:** 10.1101/2022.08.02.502485

**Authors:** Daisuke Akazawa, Hirofumi Ohashi, Takayuki Hishiki, Takeshi Morita, Shoya Iwanami, Kwang Su Kim, Yong Dam Jeong, Eun-Sil Park, Michiyo Kataoka, Kaho Shionoya, Junki Mifune, Kana Tsuchimoto, Shinjiro Ojima, Aa Haeruman Azam, Shogo Nakajima, Hyeongki Park, Tomoki Yoshikawa, Masayuki Shimojima, Kotaro Kiga, Shingo Iwami, Ken Maeda, Tadaki Suzuki, Hideki Ebihara, Yoshimasa Takahashi, Koichi Watashi

**Affiliations:** Research Center for Drug and Vaccine Development, National Institute of Infectious Diseases, Tokyo 162-8640, Japan; Interdisciplinary Biology Laboratory (iBLab), Graduate School of Science, Nagoya University, Nagoya 464-0814, Japan; Department of Science System Simulation, Pukyong National University, Busan, 48547, South Korea; Department of Mathematics, Pusan National University, Busan, South Korea; Department of Veterinary Science, National Institute of Infectious Diseases, Tokyo 162-8640, Japan; Department of Pathology, National Institute of Infectious Diseases, Tokyo 162-8640, Japan; Department of Virology II, National Institute of Infectious Diseases, Tokyo 162-8640, Japan; Department of Applied Biological Science, Tokyo University of Science, Noda 278-8510, Japan; Department of Virology I, National Institute of Infectious Diseases, Tokyo 162-8640, Japan; Institute of Mathematics for Industry, Kyushu University, Fukuoka, Japan; Institute for the Advanced Study of Human Biology (ASHBi), Kyoto University, Kyoto, Japan; Interdisciplinary Theoretical and Mathematical Sciences Program (iTHEMS), RIKEN, Saitama, Japan; NEXT-Ganken Program, Japanese Foundation for Cancer Research (JFCR), Tokyo, Japan; Science Groove Inc., Fukuoka, Japan; MIRAI, Japan Science and Technology Agency (JST), Saitama 332-0012, Japan

**Author notes:** These authors contributed equally to this work (Daisuke Akazawa, Hirofumi Ohashi). These authors contributed equally to this work (Takayuki Hishiki, Takeshi Morita). These authors contributed equally to this work (Shoya Iwanami, Kwang Su Kim, Yong Dam Jeong). Corresponding author: Koichi Watashi, Ph.D. Research Center for Drug and Vaccine Development, National Institute of Infectious Diseases, 1-23-1 Toyama, Shinjuku-ku, Tokyo 162-8640, Japan.

## Abstract

Monkeypox virus (MPXV) is a zoonotic orthopoxvirus that causes smallpox-like symptoms in humans and caused an outbreak in May 2022 that led the WHO to declare global health emergency. In this study, from a screening of approved-drug libraries using an MPXV infection cell system, atovaquone, mefloquine, and molnupiravir exhibited anti-MPXV activity, with 50% inhibitory concentrations of 0.51-5.2 μM, which is more potent than cidofovir. Whereas mefloquine was suggested to inhibit viral entry, atovaquone and molnupiravir targeted post-entry process to impair intracellular virion accumulation. Inhibitors of dihydroorotate dehydrogenase, an atovaquone’s target enzyme, showed conserved anti-MPXV activities. Combining atovaquone with tecovirimat enhanced the anti-MPXV effect of tecovirimat. Quantitative mathematical simulations predicted that atovaquone can promote viral clearance in patients by seven days at clinically relevant drug concentrations. Moreover, atovaquone and molnupiravir exhibited pan-*Orthopoxvirus* activity against vaccinia and cowpox viruses. These data suggest that atovaquone would be potential candidates for treating monkeypox.

## Introduction

Monkeypox virus (MPXV) is a zoonotic virus classified in the *Orthopoxvirus* genus of the Poxviridae family that causes smallpox-like symptoms in humans ^1, 2, 3^. After the first reports of human infection were reported in Congo in 1970, monkeypox cases have been primarily reported in Central and West African countries, with rare reports in other countries that have been linked with importation and travel from endemic African regions. In May 2022, an outbreak of human monkeypox was reported and involved more than 20,000 cases in over 70 countries mainly in Europe and North America at the time of writing of this paper ^4^. The World Health Organization (WHO) declared this monkeypox outbreak a global health emergency on July 23, 2022.

Tecovirimat and brincidofovir are approved by the US Food and Drug Administration (FDA) for the treatment of smallpox ^5^. Tecovirimat targets the viral-encoded VP37 protein to inhibit the envelopment of intracellular mature virions, whereas brincidofovir is a lipid conjugate prodrug of the nucleotide analogue, cidofovir, which inhibits viral DNA replication ^6, 7, 8^. Although these two drugs have been approved for smallpox treatment through the FDA animal efficacy rule, their effectiveness against human monkeypox has not been well documented. A recent clinical study did not show any convincing clinical benefit of brincidofovir in monkeypox patients but instead showed that the drug caused liver damage resulting in treatment cessation ^9^. Due to concerns over the international spread of monkeypox, there is increasing demand for effective and safe clinical treatments for monkeypox.

This study employed a drug repurposing approach to identify already-approved drugs that exhibit anti-MPXV activity in a virological infection assay. We also quantitatively investigated the anti-viral dynamics under clinical drug concentrations and predicted the impact on patient viral load to identify clinically relevant drug candidates. We also demonstrate the pan-*Orthopoxvirus* activity of the identified drugs and speculate on their usefulness for controlling *Orthopoxvirus*-related infectious diseases.

## Results

### Identification of approved drugs exhibiting anti-MPXV activity

Using a cell-based MPXV infection screening approach, we focused on libraries of clinically approved drugs consisting of 132 anti-viral, anti-fungal, and anti-parasitic/anti-protozoal agents (Table S1), as these classes include drugs reported to have wide range of anti-viral activity, such as remdesivir, itraconazole, and ivermectin ^10, 11, 12^. For the initial screening, we treated VeroE6 cells with MPXV at a multiplicity of infection (MOI) of 0.1 together with tested compounds, and then cytopathic effects were assessed at 72 h post-infection by microscopic observation and quantification of cell viability by high-content imaging analyzer ^13^ (see Methods for details) (Fig. 1A). The assay system was validated by observations that treatment with MPXV resulted in robust cytopathology that reduced cell viability to <1% (Fig. 1B panel b and S1A) and observations that cells were protected by treatment with a known MPXV inhibitor, tecovirimat, as a positive control ^13^ (Fig. 1B panel c and S1A). In this assay, an increase in cell survival would suggest that the tested drug inhibits virus infection/replication without cytotoxicity. In a screening at a concentration of 10 μM, 21 drugs showed > 20-fold higher cell viability than DMSO-treated controls (Fig. S1A, above the red line). As candidates, we focused on atovaquone (*anti-Pneumocystis jiroveci*), mefloquine (anti-malarial), and molnupiravir [anti-severe acute respiratory syndrome coronavirus 2 (SARS-CoV-2)] as orally-applicable drugs that are currently available clinically and exhibited remarkable anti-MPXV activity at 3.3 μM in the second screening (Fig. S1B) (see Supplementary Note-1 for details). These drugs protected cells from MPXV-induced cytopathic effects at 72 h post-infection (Fig. 1B panel d, e, and f).

**Fig. 1.**
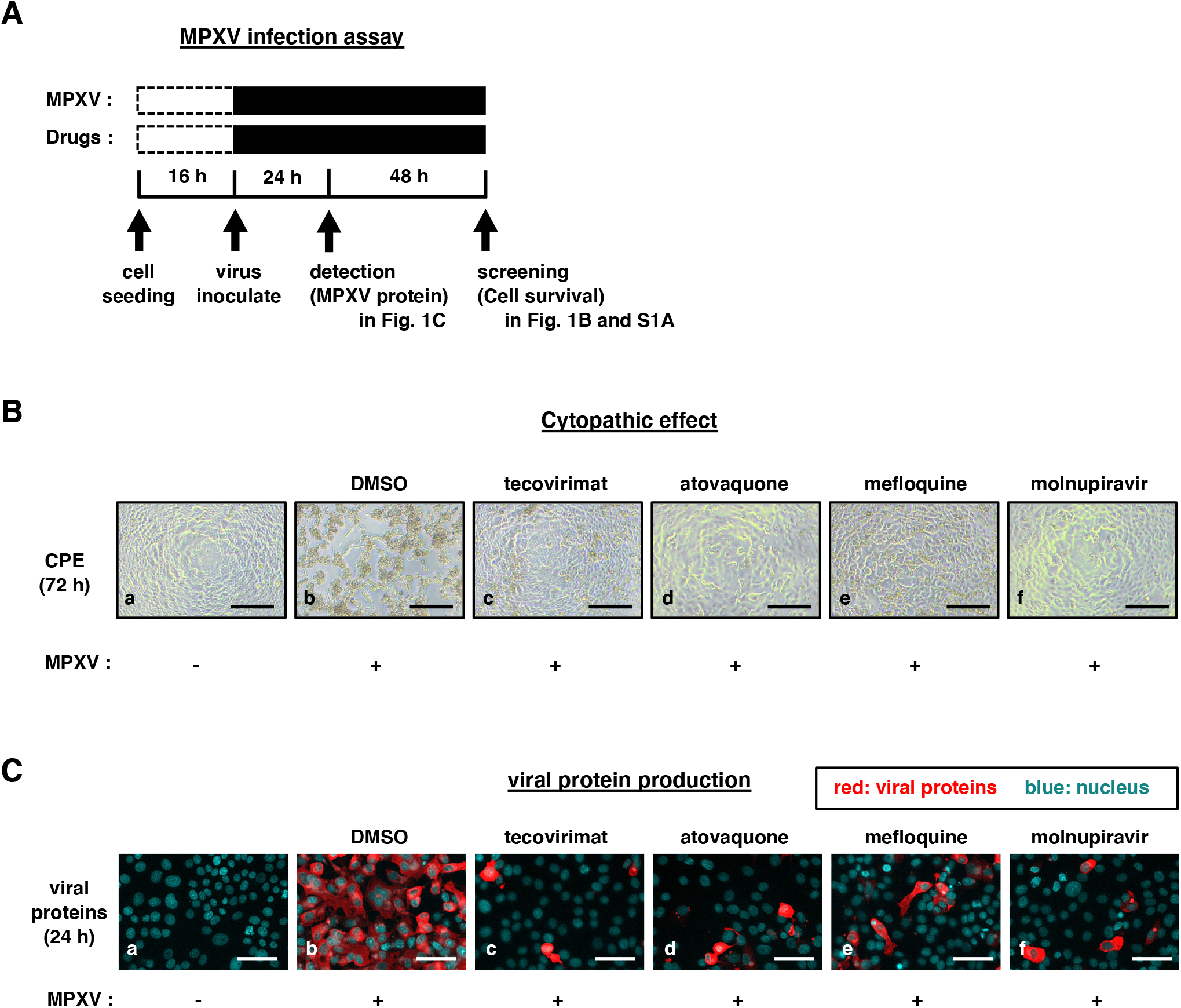
Anti-monkeypox virus (MPXV) activity of atovaquone, mefloquine, and molnupiravir. (A) Schematic representation of the MPXV infection assay. VeroE6 cells were treated with MPXV at an MOI of 0.1 and with or without drugs for 24 (Fig. 1C) or 72 h (Fig. 1B and S1A). In the screening, survived cell numbers at 72 h were measured with a high-content imaging analyzer (Fig. S1A). Cytopathic effect at 72 h was also observed by microscopy (Fig. 1B). Viral proteins in the cells were detected by immunofluorescence analysis at 24 h post-inoculation (Fig. 1C). (B) Morphology of MPXV-infected cells upon drug treatment (0.1% DMSO, 10 μM tecovirimat, 10 μM atovaquone, 10 μM mefloquine, 10 μM molnupiravir for b-f) was observed at 72 h post-inoculation by microscopy. Uninfected cells were also observed as a negative control (a). Scale bar, 200 μm. (C) MPXV protein production was detected at 24 h post-inoculation by immunofluorescence in uninfected (a) or infected VeroE6 cells upon treatment with drugs (0.1% DMSO, 5 μM tecovirimat, 5 μM atovaquone, 5 μM mefloquine, 5 μM molnupiravir for b-f). Scale bars, 50 μm. Red, MPXV proteins; Blue, nuclei.

To validate the anti-MPXV effect of these drugs, we treated VeroE6 cells with MPXV using the same protocol and detected intracellular MPXV proteins at 24 h post-infection (before sign of cytopathology emerged). As shown in Fig. 1C, these drugs dramatically reduced the production of MPXV proteins in the infection assay (Fig. 1C panel d, e, and f). These results confirmed the anti-MPXV activity of atovaquone, mefloquine, and molnupiravir.

### Anti-MPXV activity dose-response curves for atovaquone, mefloquine, and molnupiravir

The anti-MPXV activity of atovaquone, mefloquine, and molnupiravir (Fig. 2A) was assessed by quantifying viral DNA in infected cells following 30 h of treatment (before onset of MPXV-induced cytopathic effects) with the drugs at varying concentrations (Fig. 2B). We also examined the reported MPXV inhibitors tecovirimat and cidofovir ^14, 15, 16^ as positive controls. Cell viability was simultaneously quantified at different drug concentrations to examine any possible cytotoxic effects of the drugs (Fig. 2C). As shown in Fig. 2B and C, atovaquone, mefloquine, and molnupiravir reduced intracellular MPXV DNA levels in a dosedependent manner without inducing cytotoxic effects (Fig. 2B and C). The 50% and 90% maximal virus inhibitory concentrations (IC_50_ and IC_90_, summarized in Table S2) and 50% maximal cytotoxic concentrations (CC_50_) are shown in Fig. 2B and C. These three drugs exhibited greater anti-MPXV potency (lower IC_50_ and IC_90_ values) than cidofovir. In particular, the calculated IC_50_ values of atovaquone and molnupiravir were lower than the reported maximum drug concentration (C_max_) in treated patients (Table S3), suggesting the possibility that the anti-viral potency of these drugs is clinically relevant.

**Fig. 2.**
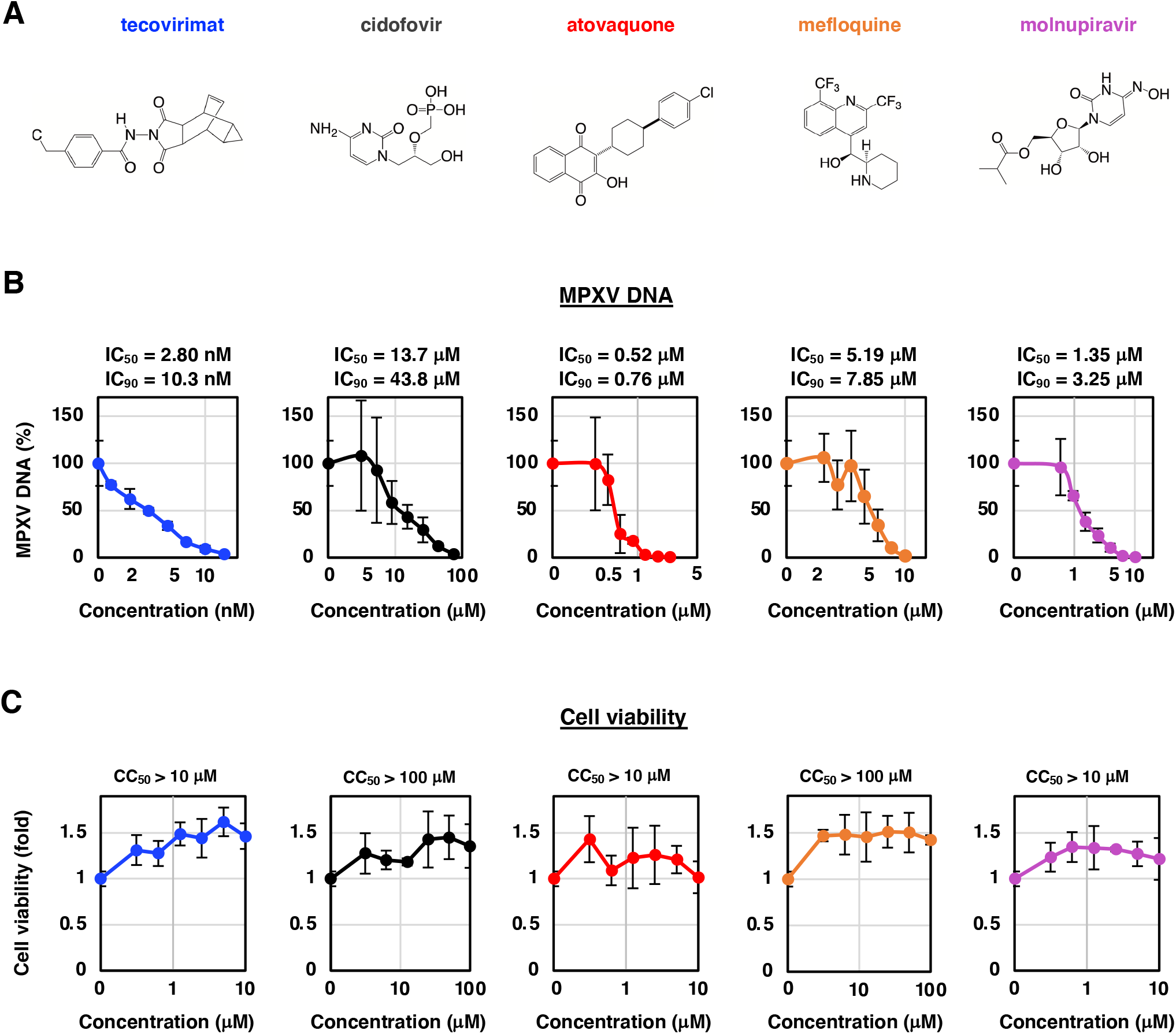
Dose-response curves for anti-MPXV activity and cytotoxicity of the drugs. (A) Chemical structures of the drugs. (B) VeroE6 cells were infected with MPXV at an MOI of 0.03 for 1 h and were washed out, followed by the incubation upon treatment with varying concentrations of each drug for another 29 h. Anti-MPXV activity was examined by quantifying MPXV DNA in cells. Relative MPXV DNA levels, determined by setting that in DMSO-treated control cells as 100%, are shown against the drug concentration (x-axis, log scale). The 50% and 90% maximal inhibitory concenstrations (IC_50_ and IC_90_) are indicated above the graphs. (C) Cell viability was also measured upon treatment with drugs at the indicated concentrations using the WST assay. The calculated 50% maximal cytotoxic concentrations (CC_50_) are shown above the graphs.

### Viral life cycle step targeted by the identified drugs

In the viral life cycle, MPXV attaches to target cells and enter inside cells to deliver the viral core into the cytoplasm (entry phase, Fig. 3A). After early transcription, protein synthesis, and core uncoating, the viral DNA replicates as well as drives intermediate and late transcriptions, and then assembled into virions in the cytoplasmic viral factories, followed by a stepwise virion maturation process resulting in the production of progeny infectious virions (post-entry phase, Fig. 3A) ^17^. To determine which phase in the viral life cycle is inhibited by the identified drugs, we performed time-of-addition assay in which cells were treated with the drugs at varying time points (Fig. 3B, left) to distinguish the entry and post-entry phases ^18, 19^. Drug were administered either (a) throughout the assay (whole life cycle), (b) for the initial 2 h (entry phase), or (c) for the last 22 h after viral infection (post-entry and re-infection phase) (Fig. 3B, left). The positive control tecovirimat, which inhibits virion maturation, exhibited significant anti-viral activity under conditions (a) and (c) but not (b), whereas heparin, which reportedly inhibits viral entry ^20^, showed significant anti-MPXV activity in condition (b) (Fig. 3B, right), thereby validating the time-of-addition assay. In this assay, mefloquine showed significant anti-MPXV activity under condition (b), in addition to conditions (a) and (c) (Fig. 3B, right), consistent with the reported inhibition of the entry of multiple viruses, including coronaviruses and Ebola virus ^19, 21^, although there are no reports of its effect on poxviruses. In contrast, atovaquone and molnupiravir predominantly reduced MPXV DNA levels in cells treated under condition (c) but not condition (b) (Fig. 3B, right), suggesting that atovaquone and molnupiravir target the post-entry phase.

**Fig. 3.**
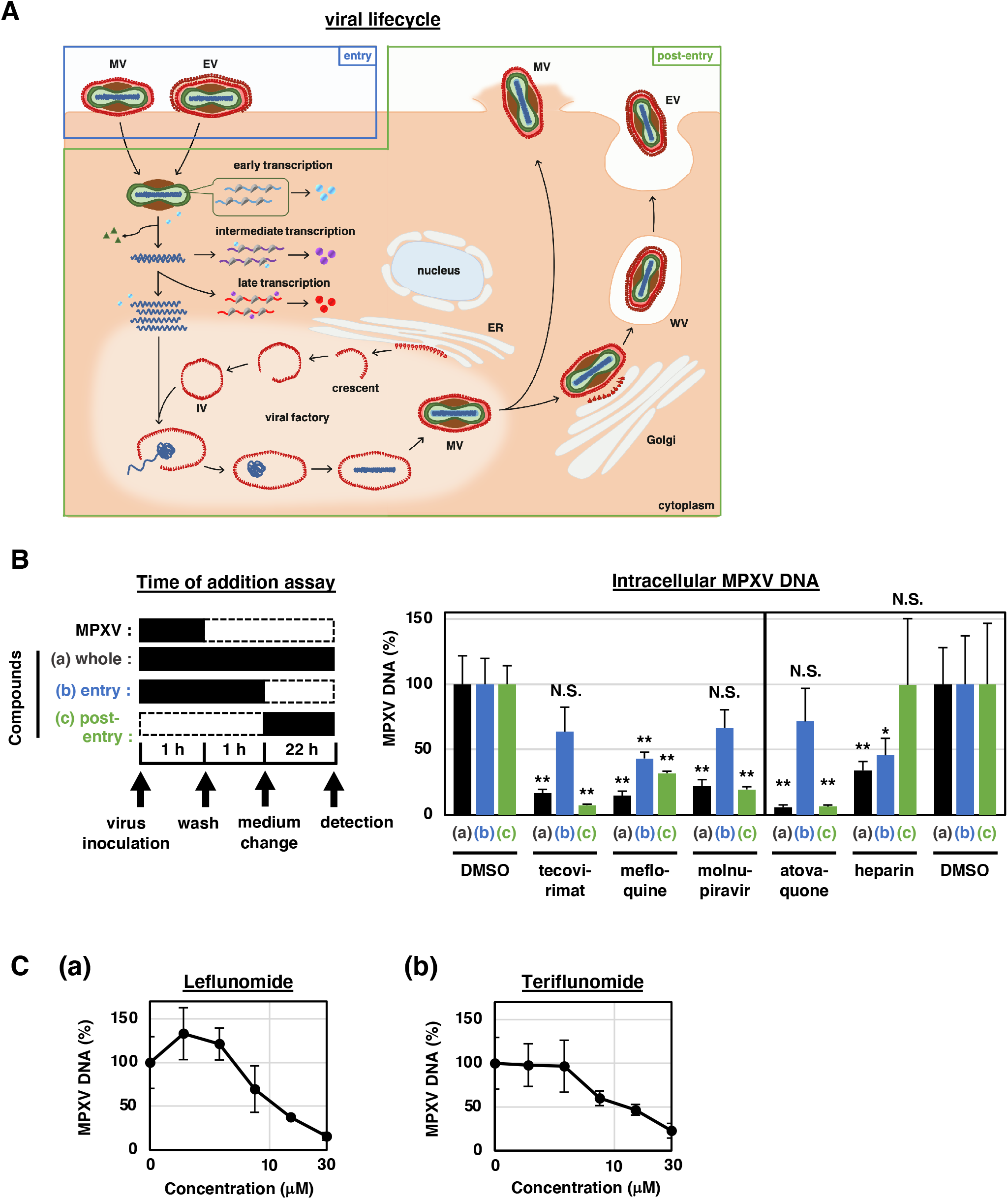
Viral life cycle step targeted by drugs. (A) Schematic illustration of the MPXV life cycle. Target steps of the positive control compounds, heparin and tecovirimat are also shown. (B) Time-of-addition assay. Left, experimental schedules of the assays in which cells were treated with drugs at three different times, either (a) whole: throughout the assay for 24 h, (b) entry: for the initial 2 h (1 h with the MPXV inoculum and the following 1 h after inoculation), or (c) post-entry: for the last 22 h after inoculation. Right, MPXV DNA levels in infected VeroE6 cells treated with the drugs under conditions (a), (b), or (c) were determined and are shown relative to that of the DMSO-treated control. (C) Dose-dependent anti-MPXV activity of leflunomide and teriflunomide, DHODH inhibitors. Anti-MPXV activities were examined as shown in Fig. 2B at the indicated concentrations.

Molnupiravir is a nucleoside analogue that targets polymerization of the genome of different virus classes ^22, 23, 24, 25^, consistent with the observed inhibition of MPXV replication. Atovaquone targets the cytochrome bc1 complex to inhibit mitochondrial electron transport ^26^ and also inhibits parasite dihydroorotate dehydrogenase (DHODH) in the pyrimidine biosynthesis pathway ^27^, which regulates the replication of a wide range of viruses including influenza viruses, coronaviruses, and flaviviruses ^28^. We showed that other DHODH inhibitors, leflunomide and teriflunomide ^29^, also reduced intracellular MPXV DNA levels in a dose-dependent manner (Fig. 3C). These data suggest that the inactivation of DHODH inhibited MPXV replication.

### Electron microscopic analysis of drug-treated infected cells

Infection of cells with MPXV induces the formation of cytoplasmic viral factories to initiate the assembly of virions that sequentially forms crescents (Cs), immature virions (IVs), and mature virions (MVs), which further follow the maturation process to wrapped virions (WVs) and extracellular virions (EVs) ^30^. To examine the effect of the identified drugs on virion maturation and the formation of cytoplasmic viral factories, we used transmission electron microscopy to examine the intracellular morphology of infected drug-treated cells, including cytoplasmic viral factories and the virions produced. As shown in Fig. 4, the cytoplasm of infected cells exhibited regions with high electron density at perinuclear areas, indicative of cytoplasmic viral factories (Fig. 4 panel b, *) ^30^. In contrast, no such regions were observed in uninfected cells (Fig. 4 panel-a). These factories contained Cs, circular IVs, and dense MVs (Fig. 4 panel b’). WVs were also observed outside of the factories (Fig. 4 panel b”). Upon tecovirimat treatment, a significant number of cells exhibited accumulation of Cs, IVs and MVs at the cytoplasmic viral factories (Fig. 4 panel c, * and c’), confirming that this drug inhibits the maturation to WVs but not viral replication before virion production. In contrast, treatment with atovaquone and molnupiravir resulted in a dramatic reduction in the number of infected cells, and few virions were observed in these cells, even though they possessed cytoplasmic viral factories (Fig. 4 panel d, e, *). In contrast to tecovirimat, these observations suggest that atovaquone and molnupiravir inhibit MPXV replication before virion maturation.

**Fig. 4.**
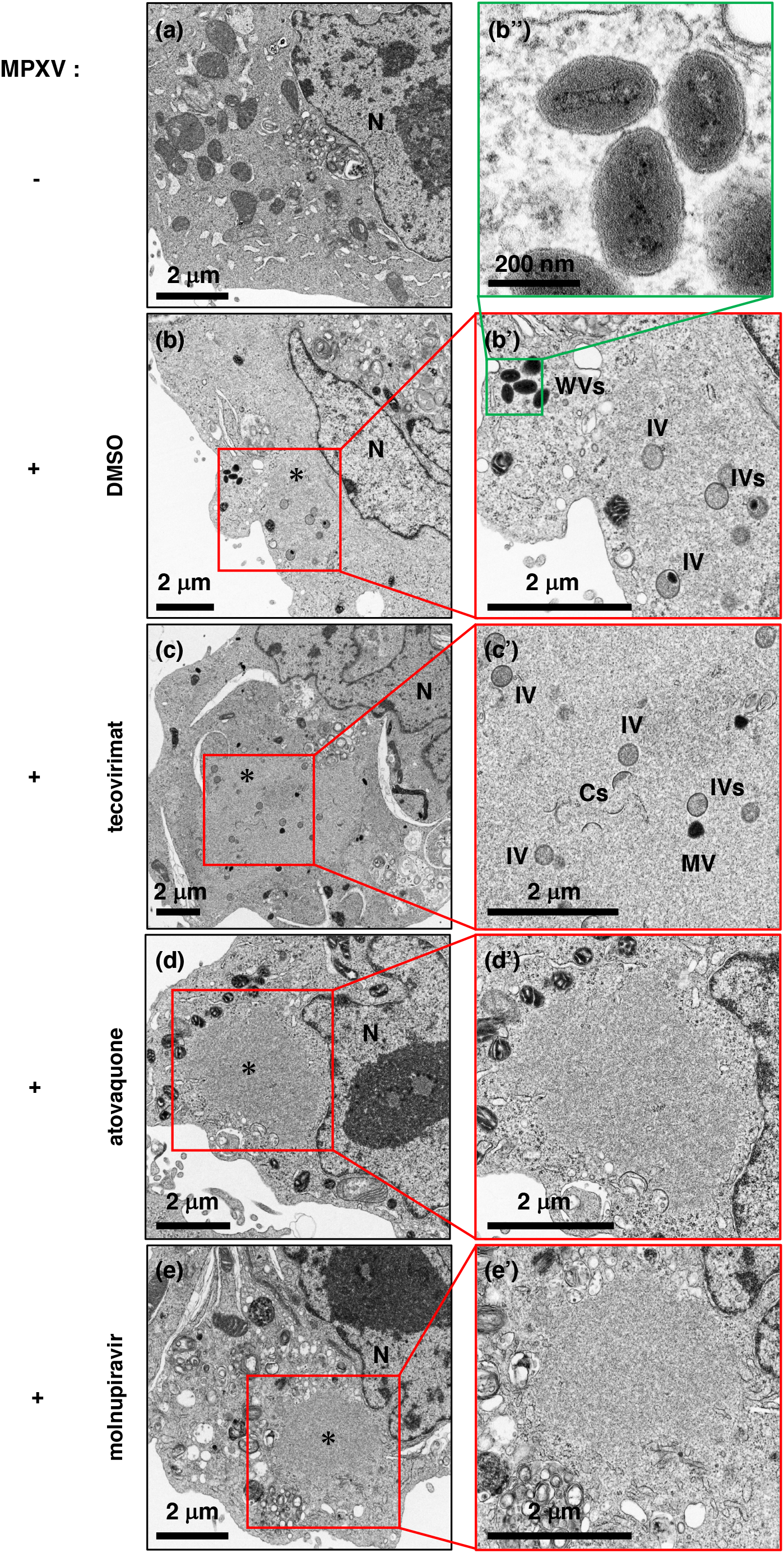
Atovaquone and molnupiravir reduced virion production in MPXV-infected cells. VeroE6 cells infected with (b-e) or without (a) MPXV at an MOI of 0.1 were incubated upon drug treatment (b, 0.1% DMSO; c, 5 μM tecovirimat; d, 5 μM atovaquone; e, 5 μM molnupiravir) for 24 h. The cells were trypsinized and observed by transmission electron microscopy. A total of 150 cells were observed for each sample, and the figure shows the representative images of infected cell morphology. The images in b’, c’, d’, and e’ show the boxed areas in b, c, d, and e, respectively, at higher magnification. Panel b’’ is a higher-magnification image of the inset in b’ with 90° rotation. Scale bars in a, b, b’, c, c’, d, d’, e, and e’, 2 μm; that in b’’, 200 nm. N, nucleus; *, cytoplasmic viral factories; C, crescent; IV, immature virion; MV, mature virion; WV, wrapped virion.

### Impact of anti-viral drugs on MPXV infection in clinical settings

Combining the pharmacokinetics (PK) information (summarized in Table S3) of these approved drugs when administered in patients with the observed dosedependent anti-viral MPXV activity information [pharmacodynamics (PD) information: summarized in Table S2], we calculated the anti-viral effect for clinical drug concentrations using the PK/ PD model (Eq.(11) in Supporting Note-2) (Fig. 5A). We assumed a simple one-compartment model ^18^ with the reported maximum drug concentrations in plasma (C_max_) and half-life values of tecovirimat ^31^, cidofovir ^32^, atovaquone ^33^, mefloquine ^34^, and molnupiravir ^35^ by administration at approval doses (see Table S3). We found that cidofovir and molnupiravir show anti-viral effects after administration but which are rapidly declined, reflected by their short half-lives, whereas the anti-viral effects of tecovirimat and atovaquone are maintained at high levels during drug treatment (Fig. 5A). That of mefloquine was estimated to gradually decline after administration (Fig. 5A).

**Fig. 5.**
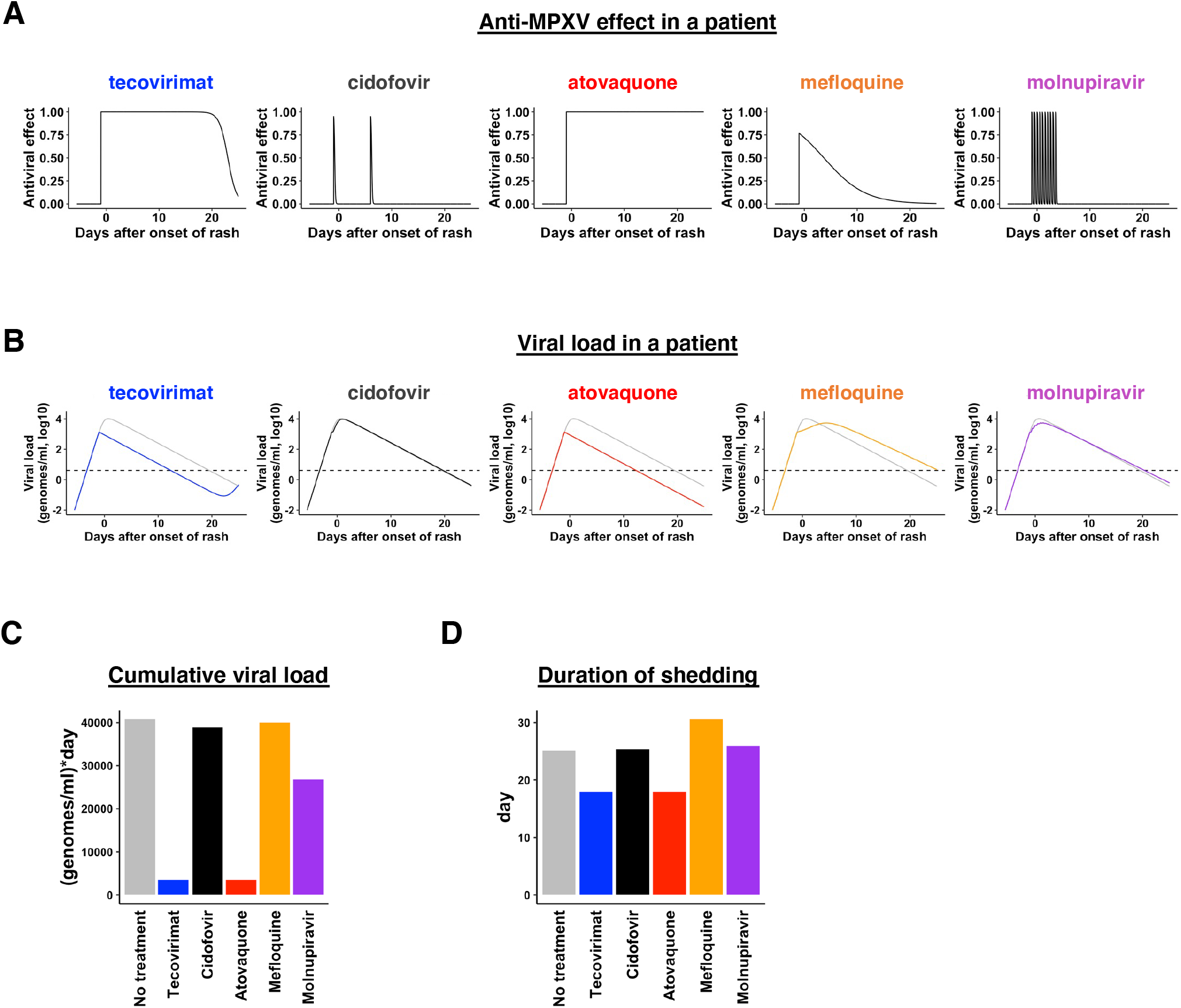
Mathematical prediction of the impact of the identified drugs on viral load dynamics in clinical settings. (A) Time-dependent antiviral effects of tecovirimat (600 mg, oral BID for 14 days), cidofovir (5 mg/kg, intravenous qWeek × 2 doses), atovaquone (750 mg, oral BID for 21 days), mefloquine (25 mg/kg, oral once) and molnupiravir (800 mg, oral BID for 5 days) predicted by PK/PD modeling are shown. (B) Viral infection dynamics in the presence or absence of tecovirimat, cidofovir, atovaquone, mefloquine, and molnupiravir by PK/PD/ VD models are shown. The gray lines show the predicted viral load in patients in the absence of treatments. The colored lines show the expected viral load in the presence of treatments. The dashed lines indicate the detection limit of MPXV DNA. (C, D) Cumulative viral load [area under the curve in (B)] and the duration of virus shedding [time until the viral load is below the detection limit in (B)] were calculated for untreated (gray) and treated (colored) patients.

To evaluate the impact of drug treatment on MPXV-infected patients after onset of rash in patients (i.e., day 0 is the date of onset of rash), we developed a mathematical model [i.e., PK/PD/viral dynamics (VD) model: Eqs.(7-11) in Supporting Note-2] integrating the PK/PD information with VD information (summarized in Table S4) (Fig. S2). As shown in Fig. 5B, in the absence of antiviral treatment, the mathematical model [corresponding to Eqs.(3-4) in Supporting Note-2] predicted that MPXV viral load would exponentially increases for the first 0.71 days and then peak, which would be followed by a gradual decline (Fig. 5B, gray, Fig. S3). Based on the expected viral load, we calculated the cumulative viral RNA burden (i.e., area under the curve of viral load) (Fig. 5C and Fig. S2) and the duration of viral shedding (Fig. 5D and Fig. S2). Based on the time-dependent antiviral effect of the drugs, we predicted the impact of the anti-viral effects on the dynamics of MPXV infection when the drugs are administrated on −1 days, which corresponds to the date for the first viral load sampling, after the onset of rash (Fig. 5B, colored). Interestingly, our quantitative simulations predicted that atovaquone would be the most effective in the clinical setting, reducing the cumulative viral load by 91.6% (Fig. 5C) and the duration of virus positivity in serum by 7.16 days shorter duration compared with untreated control subjects (Fig. 5D). As we explored in the recent study ^36^, it is required for reductions of viral shedding that antiviral treatments start before the viral load hits its peak.

We here used the previously reported mean values of the viral load (genomes/ml) in the blood ^37^ for parameter estimations for our quantitative simulation (Fig. S3 and Supporting Note-2). It should be noted that the peak PCR viral load in blood occurs near the first day of rash appearance, meaning viral load may have already peaked. A time-course individual-level clinical viral load from different specimens covering whole MPXV infection is required to accurately evaluate the effect of antivirals, although clinical viral load data is quite limited so far.

### Combined treatment with atovaquone and tecovirimat

The clinical outcome of an anti-viral treatment regimen can be improved by combining drugs, as is clinically employed in the treatment of human immunodeficiency virus and hepatitis C virus infections ^38, 39^. Based on possible differences in the mode of action of atovaquone and tecovirimat, we examined the anti-viral activity of the combination of these drugs using the MPXV infection assay. Cells were treated with combinations of the drugs at various concentrations for 30 h, after which intracellular viral DNA and cell viability were evaluated. Compared with the dose-dependent reduction in viral DNA levels observed with single treatment with atovaquone or tecovirimat, the combination of these drugs further reduced viral DNA levels (Fig. 6A). We did not observe any significant cytotoxicity at any of the drug concentrations tested (Fig. 6B). We then compared the observed experimental anti-viral activity with theoretical predictions calculated using a classical Bliss independence model that assumes the drugs act independently (Supporting Note-2) ^18, 40, 41^. The difference between the observed values and theoretical predictions suggests that atovaquone and tecovirimat exhibit a synergistic activity over a broad range of concentrations, especially at lower concentration ranges (Fig. 6C red: synergistic effect) (High concentrations readily showed enough activities by single treatment, which appears to be calculated as less synergistic effects). Thus, combination treatment with atovaquone enhances the anti-viral activity of tecovirimat.

**Fig. 6.**
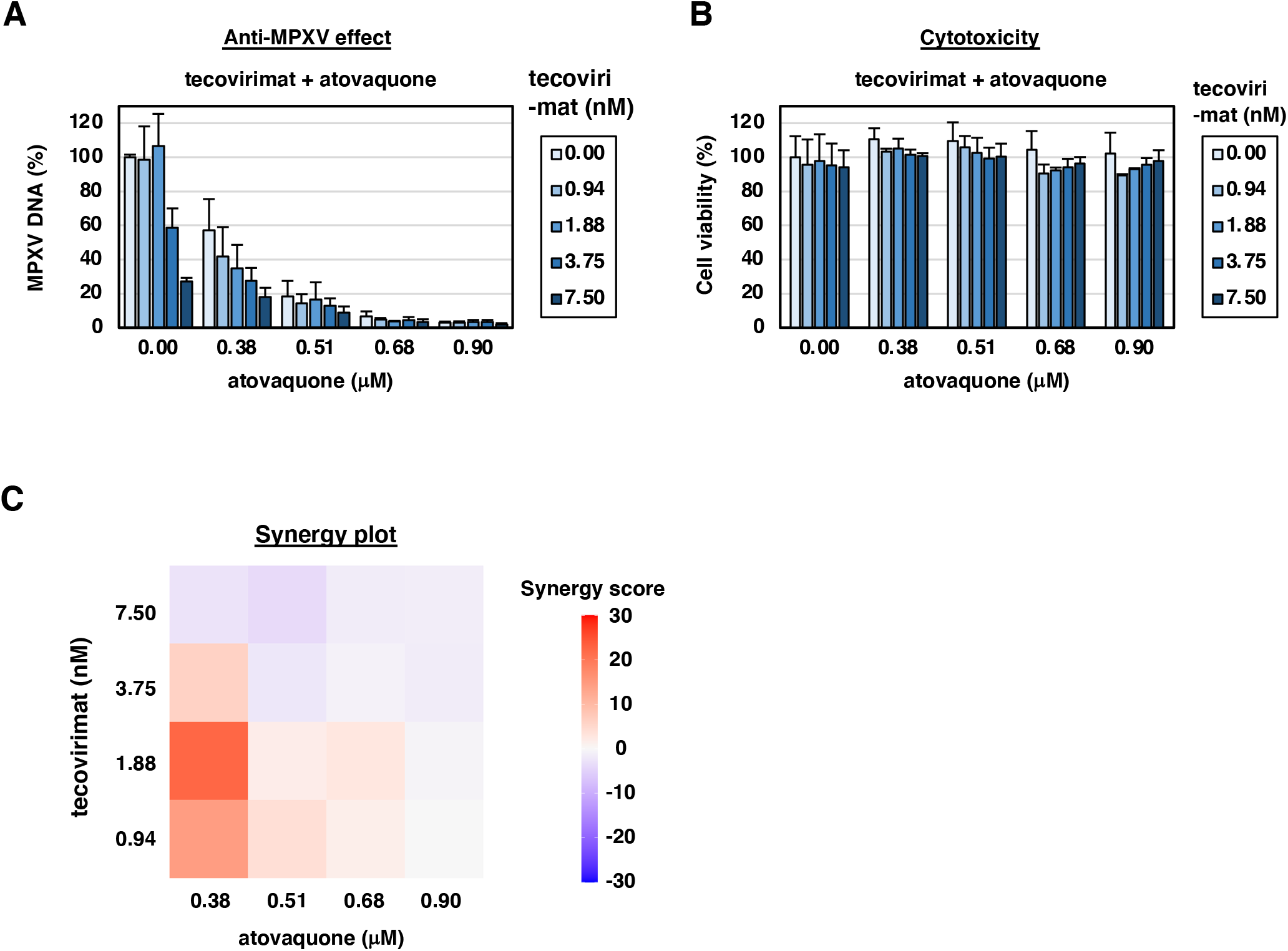
Cotreatment with tecovirimat and atovaquone. (A) Viral DNA in VeroE6 cells cotreated with tecovirimat and atovaquone at varying concentrations [tecovirimat: 0, 0.94, 1.88, 3.75, and 7.5 nM (x 2 dilution), atovaquone: 0, 0.38, 0.51, 0.68, and 0.90 μM (x 1.3 dilution)] for 30 h was quantified and are shown relative to the DMSO-treated control. (B) Cell viability was also measured at 30 h post-treatment. (C) Heatmap of synergy scores for tecovirimat and atovaquone is shown based on a Bliss independence model. Red, white, and blue colors on the heatmap indicate the synergistic, additive, and antagonistic interactions between the two drugs, respectively.

### Pan-anti-*Orthopoxvirus* activity of atovaquone and molnupiravir

Pan-anti-*Orthopoxvirus* drugs are urgently needed as a means of combating future poxvirus outbreaks or possible uses of such viruses in bioterrorism ^5^. Our study demonstrated that atovaquone, mefloquine, and molnupiravir exhibited anti-viral activity against the MPXV Liberia strain (Fig. 7A) as well as the Zr599 strain of MPXV, as shown by immunofluorescence analysis (Fig. 1). In addition, these drugs reduced the expression of viral proteins in infection assays involving other orthopoxviruses, vaccinia virus and cowpox virus, except for no significant reduction of vaccinia virus proteins by mefloquine (Fig. 7B and C). These results suggest that atovaquone and molnupiravir exhibit a pan-anti-*Orthpoxvirus* potential.

**Fig. 7.**
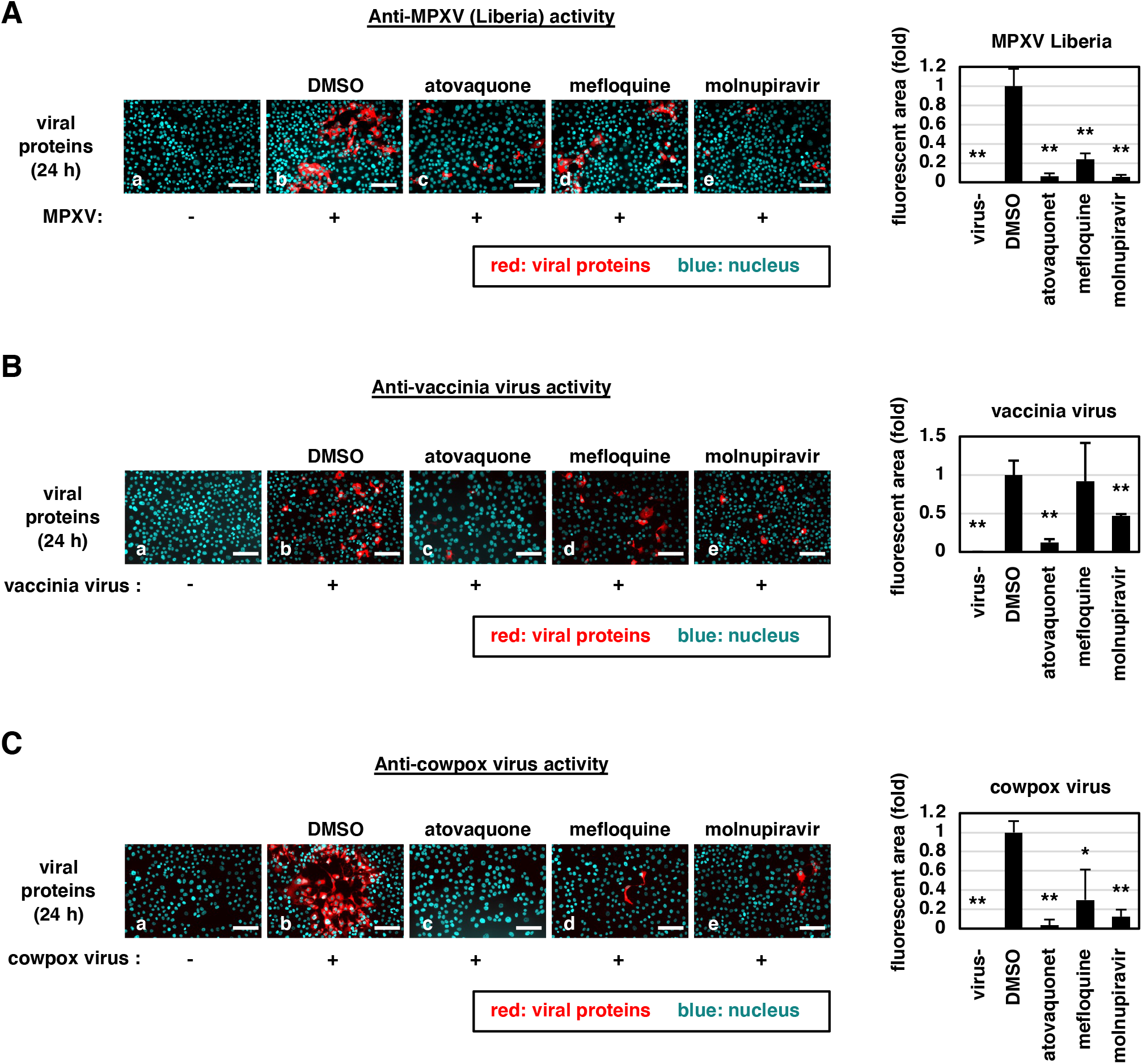
Anti-viral effect of atovaquone, mefloquine, and molnupiravir against orthopoxviruses. VeroE6 cells were inoculated with (b-e) or without(a) MPXV (Liberia strain) (A), vaccinia virus (B), or cowpox virus (C) and then treated with drugs (0.1% DMSO, 5 μM atovaquone, 5 μM mefloquine, 5 μM molnupiravir for b-e) for 24 h. Viral proteins (red) as well as the nuclei (blue) were detected by immunofluorescence. Scale bars, 100 μm. Observed areas of red fluorescence were quantified and are shown relative to the DMSO-treated control (right graphs).

## Discussion

In this study, we screened a library of approved drugs for anti-MPXV activity using a cell culture infection assay and identified atovaquone, mefloquine, and molnupiravir, as candidate drugs. The IC_50_ of atovaquone was 0.516 μM, which is within the concentration range for clinical use, with 31.3 μM as the plasma C_max_ and 67.0 h as the half-life ^33^. Our mathematical modeling predicted that atovaquone would exhibit sustained anti-MPXV activity following administration at approved doses and induce rapid viral decay in infected patients, reducing the cumulative viral load and shortening the time until virus elimination. Another potential application for atovaquone is combination use with approved anti-*Orthopoxvirus* agents such as tecovirimat. Interestingly, addition of atovaquone to tecovirimat in cell culture experiments resulted in a further reduction in MPXV DNA levels without cytotoxic effects. Given the good tolerability profile of atovaquone in clinical settings ^42^, our present data provide attractive idea for supplementing atovaquone to improve the current approved anti-orthopoxvirus treatment.

Mechanistically, molnupiravir is a nucleoside analogue that targets the polymerization of the genome of a variety of different viruses, including hepatitis C virus, norovirus, chikungunya virus, and coronaviruses ^22, 23, 24, 25^. It is speculated that molnupiravir also targets polymerization of the MPXV genome, and this should be demonstrated in future studies. Mefloquine reportedly inhibits the cell entry of multiple viruses, including coronaviruses and Ebola virus, although the mode of action remains unclear ^19, 21^. Our analysis of atovaquone also suggests that it affects DHODH in MPXV replication. Indeed, the DHODH inhibitor brequinar reportedly inhibits the replication of other poxviruses, Cantagolo virus, vaccinia virus, and cowpox virus ^43^, supporting our proposed mechanism. Clinical C_max_ and half-life for teriflunomide, an active metabolite of a rheumatoid arthritis agent, leflunomide, by administration of leflunomide at 5-20 mg daily is reportedly 107-192 μM and 2 weeks ^44^, suggesting another candidate for an anti-MPXV agent. Also, searches for new DHODH inhibitors should enable the development of more potent anti-MPXV agents in the future.

The anti-viral activities of atovaquone against MPXV, vaccinia virus, and cowpox virus, were reproduced in multiple cell lines (Fig. S4). But, limitation of our study was that it used only a cell culture infection assay without animal model experiments and patient studies. However, considering the limited amount of research regarding this virus to date and the current outbreak and spread of monkeypox across multiple continents, we believe rapid analyses using cell culture infection assays will be of benefit in providing scientific evidences to propose alternative treatment options for this infectious disease and to minimize its international spread. It should also be highlighted that these drugs exhibit anti-viral activity against multiple orthopoxviruses, making them as candidates for controlling a wide range of orthopoxviruses, potential zoonotic and bioterrorism agents. Further studies on animals and human subjects in the future should lead to the development of alternative and/or better treatments.

## Methods

### Cell culture

VeroE6 cells were cultured in Dulbecco’s modified Eagle’s medium (DMEM; FUJIFILM Wako) supplemented with 5% fetal bovine serum (FBS; NICHIREI), and 100 units/mL penicillin-streptomycin (Thermo Fisher Scientific) at 37°C in 5% CO_2_ ^19^. During infection assays, the concentration of fetal bovine serum in the medium was changed from 5% to 2%. Huh-7 cells were supplemented with DMEM supplemented with 10% FBS, 100 units/mL penicillin-streptomycin, 0.1 mM nonessential amino acids (Invitrogen), 1 mM sodium pyruvate, and 10 mM HEPES (pH 7.4) at 37 °C in 5% CO2 ^19^.

### Infection assay

MPXV was handled in a biosafety level 3 (BSL3) facility. MPXV stocks of the Zr-599 and Liberia strains were prepared by propagating viruses in VeroE6 cells; virus infectious titers were determined by plaque assay ^13^. The data shown in Fig. 1, 2, 3, 6, and S1 were generated using the Zr599 strain, and the data in Fig. 4 and 7A were generated using the Liberia strain. For infection assays, VeroE6 (Fig. 1, 2, 3, 4, 6, 7, S1) or Huh-7 cells (Fig. S4) were inoculated with MPXV at a multiplicity of infection (MOI) of 0.1 (Fig. 1B, 4, 7A, and S1) or 0.03 (Fig. 1C, 2, and 6). The cells were cultured with the virus inoculum for either 72 h (Fig. 1B and S1A), 24 h (Fig. 1C and 4), or 1 h followed by washing out and incubating with fresh virus-free medium for an additional 29 h (Fig. 2 and 6) or 23 h (Fig. 3).

Vaccinia virus (LC16m8) and cowpox virus (Brighton Red) were inoculated to VeroE6 cells at an MOI of 0.1 upon treatment with each drug for 24 h to detect viral proteins by immunofluorescence as described below ^45, 46^ (Fig. 7B and C). Viruses at the same amount were inoculated to Huh-7 cells and virus proteins were detected with the same protocol (Fig. S4).

### Drug library and screening

The drug library used for screening included 65 anti-viral, 33 anti-fungal, and 34 anti-protozoal/anti-parasitic approved drugs (Selleck), as shown in Table S1. For drug screening, VeroE6 cells were seeded in 96 well plates and inoculated with MPXV at an MOI of 0.1 together with treatment with each drug at 10 μM (8 drugs treated at 2 μM are shown in Table S1) for 72 h, as shown in Fig. 1A. The cells were recovered by fixing in 4% paraformaldehyde and then were treated with 0.02% 4’,6-diamidino-2-phenylindole (DAPI) for nuclear staining. The number of surviving cells was determined using a high-content imaging analyzer ImageXpress Micro Confocal (MOLECULAR DEVICES), as described previously ^18^ (Fig. S1A). Drugs that augmented the number of surviving cells upon MPXV infection by more than 20-fold relative to the DMSO-treated control were selected as first hits. Among the first hits, the drugs ultimately chosen for focus in this study were further selected according to the criteria shown in Supplementary Note-1.

### Immunofluorescence analysis

Viral proteins in cells infected with MPXV, vaccinia virus, or cowpox virus were detected by indirect immunofluorescence analysis, essentially as described ^18^, using a rabbit anti-vaccinia virus antibody (Abcam) as the primary antibody and anti-rabbit Alexa Fluor Plus 555 (Thermo Fisher Scientific) as the secondary antibody (Fig. 1C, 7, and S4). Nuclei were stained with DAPI. Fluorescence images were acquired using a fluorescent microscope (BZ-X710; Keyence). Red and blue signals in figures show viral proteins and cell nuclei, respectively. For quantification, the area of red fluorescence was determined by Dynamic Cell Count (Keyence) (Fig. 7).

### Real-time PCR

Intracellular DNA was extracted by using a QIAamp DNA Mini Kit (QIAGEN) according to the manufacturer’s instructions, and MPXV DNA was analyzed using real-time PCR with TaqMan Gene Expression Master Mix (Applied Biosystems) with 5’-GAGATTAGCAGACTCCAA-3’ as the forward primer and 5’-GATTCAATTTCCAGTTTGTAC-3’ as the reverse primer and 5’-FAM-GCAGTCGTTCAACTGTATTTCAAGATCTGAGAT-3’-TAMRA as the probe ^46^.

### Cell viability assay

Cell viability was measured using the WST assay (Cell Counting Kit-8; DOJINDO) according to the manufacturer’s protocol (Fig. 2C and 6B).

### Time-of-addition assay

VeroE6 cells were inoculated with MPXV at an MOI of 0.1 and incubated for 1 h, after which unbound viruses were removed by washing. The cells were cultured with compounds for three different time periods before examining the anti-viral activity of the drugs by measuring intracellular viral DNA levels (Fig. 3B) ^18, 19^. Cells were treated with the drugs either (a) throughout the entire assay for 24 h, covering the whole viral life cycle (whole); (b) only for the initial 2 h (1 h during virus inoculation + 1 h after washing to remove free viruses) to cover the viral entry step (entry); or (c) for the last 22 h, starting after virus inoculation until harvest, covering the time of post-entry and re-infection (post-entry).

### Electron microscopy

Cells were treated with or without MPXV at an MOI of 0.1 and with each drug for 24 h. The cells were trypsinized and fixed with buffer [2.5% glutaraldehyde, 2% PFA, and 0.1 M phosphate buffer (pH7.4)] for 4 days at 4°C, followed by postfixation with 1% osmium tetroxide, staining with 0.5% uranyl acetate, dehydration with graded series of alcohols, and embedding with epoxy resin ^47^. Ultrathin sections were stained with uranyl acetate and lead citrate and observed under a transmission electron microscope. At least 150 cells per each sample in the ultrathin sections were observed, and representative images are showed in Fig. 4.

### Mathematical analysis

Quantification of dose-response relationships of drugs, determination of synergism between atovaquone and tecovirimat, prediction of the anti-MPXV effect of the drugs, and drug impact on MPXV infection are shown in detail in Supporting Note-2.

### Drug cotreatment

For the results shown in Fig. 6, VeroE6 cells were inoculated with MPXV for 1 h and were then washed out, followed by incubation for an additional 29 h. The cells were then incubated with the drugs singly or in combination at different concentrations [tecovirimat: 0, 0.94, 1.88, 3.75, and 7.5 nM (x 2 dilution), atovaquone: 0, 0.38, 0.51, 0.68, and 0.90 μM (x 1.3 dilution)] for 1 h during virus inoculation and 29 h after inoculation. Intracellular MPXV DNA was quantified to assess anti-viral activity. Cell viability was also measured to examine the cytotoxicity of the drugs.

### Statistical analyses

The statistical significance was analyzed by using the two-tailed Student’s *t*-test (*p < 0.05; **p < 0.01; N.S., not significant).

## Acknowledgments

This work was supported by The Agency for Medical Research and Development (AMED) (JP21fk0108589, JP22fk0310504, JP22jm0210068, JP20wm0325007), the Japan Society for the Promotion of Science KAKENHI (JP20H03499), the JST MIRAI program (JPMJMI22G1), Moonshot R&D (JPMJMS2021 and JPMJMS2025), and the Takeda Science Foundation.

## Author Contributions

D.A., H.O., T.M., T.H., E.S.P., M.K., K.S., J.M., K.T., S.O., A.H.A., S.N., K.K., K.W., performed experiments. Y.D.J., S.Iwanami, K.S.K., S.Iwami, analyzed mathematical data. T.Y., M.S., K.M., T.S., H.E., Y.T., contributed materials. K.S., J.M., S.Iwami, K.W., wrote paper. S.Iwami, K.W., acquired funding. K.W., supervised project.

## References

1. Kozlov M. Monkeypox goes global: why scientists are on alert. Nature 606, 15–16 (2022).

2. Thornhill JP, et al. Monkeypox Virus Infection in Humans across 16 Countries - April-June 2022. N Engl J Med, (2022).

3. Titanji BK. Neglecting emerging diseases - monkeypox is the latest price of a costly default. Med (N Y) 3, 433–434 (2022).

4. CDC. 2022 Monkeypox Outbreak Global Map. https://wwwcdcgov/poxvirus/monkeypox/response/2022/world-maphtml.

5. Delaune D, Iseni F. Drug Development against Smallpox: Present and Future. Antimicrob Agents Chemother 64, (2020).

6. Duraffour S, et al. ST-246 is a key antiviral to inhibit the viral F13L phospholipase, one of the essential proteins for orthopoxvirus wrapping. J Antimicrob Chemother 70, 1367–1380 (2015).

7. Hutson CL, et al. Pharmacokinetics and Efficacy of a Potential Smallpox Therapeutic, Brincidofovir, in a Lethal Monkeypox Virus Animal Model. mSphere 6, (2021).

8. Parker S, et al. Efficacy of therapeutic intervention with an oral ether-lipid analogue of cidofovir (CMX001) in a lethal mousepox model. Antiviral Res 77, 39–49 (2008).

9. Adler H, et al. Clinical features and management of human monkeypox: a retrospective observational study in the UK. Lancet Infect Dis 22, 1153–1162 (2022).

10. Bauer L, et al. Structure-activity relationship study of itraconazole, a broad-range inhibitor of picornavirus replication that targets oxysterol-binding protein (OSBP). Antiviral Res 156, 55–63 (2018).

11. Jans DA, Wagstaff KM. The broad spectrum host-directed agent ivermectin as an antiviral for SARS-CoV-2 ? Biochem Biophys Res Commun 538, 163–172 (2021).

12. Warren TK, et al. Therapeutic efficacy of the small molecule GS-5734 against Ebola virus in rhesus monkeys. Nature 531, 381–385 (2016).

13. Saijo M, et al. Virulence and pathophysiology of the Congo Basin and West African strains of monkeypox virus in non-human primates. J Gen Virol 90, 2266–2271 (2009).

14. Smee DF, Sidwell RW, Kefauver D, Bray M, Huggins JW. Characterization of wildtype and cidofovir-resistant strains of camelpox, cowpox, monkeypox, and vaccinia viruses. Antimicrob Agents Chemother 46, 1329–1335 (2002).

15. Stittelaar KJ, et al. Antiviral treatment is more effective than smallpox vaccination upon lethal monkeypox virus infection. Nature 439, 745–748 (2006).

16. Yang G, et al. An orally bioavailable antipoxvirus compound (ST-246) inhibits extracellular virus formation and protects mice from lethal orthopoxvirus Challenge. J Virol 79, 13139–13149 (2005).

17. Schramm B, Locker JK. Cytoplasmic organization of POXvirus DNA replication. Traffic 6, 839–846 (2005).

18. Ohashi H, et al. Potential anti-COVID-19 agents, cepharanthine and nelfinavir, and their usage for combination treatment. iScience 24, 102367 (2021).

19. Shionoya K, et al. Mefloquine, a Potent Anti-severe Acute Respiratory Syndrome-Related Coronavirus 2 (SARS-CoV-2) Drug as an Entry Inhibitor in vitro. Front Microbiol 12, 651403 (2021).

20. Vazquez MI, Esteban M. Identification of functional domains in the 14-kilodalton envelope protein (A27L) of vaccinia virus. J Virol 73, 9098–9109 (1999).

21. Sun W, et al. Synergistic drug combination effectively blocks Ebola virus infection. Antiviral Res 137, 165–172 (2017).

22. Costantini VP, et al. Antiviral activity of nucleoside analogues against norovirus. Antivir Ther 17, 981–991 (2012).

23. Ehteshami M, et al. Characterization of beta-d-N(4)-Hydroxycytidine as a Novel Inhibitor of Chikungunya Virus. Antimicrob Agents Chemother 61, (2017).

24. Kabinger F, et al. Mechanism of molnupiravir-induced SARS-CoV-2 mutagenesis. Nat Struct Mol Biol 28, 740–746 (2021).

25. Stuyver LJ, et al. Ribonucleoside analogue that blocks replication of bovine viral diarrhea and hepatitis C viruses in culture. Antimicrob Agents Chemother 47, 244–254 (2003).

26. Birth D, Kao WC, Hunte C. Structural analysis of atovaquone-inhibited cytochrome bc1 complex reveals the molecular basis of antimalarial drug action. Nat Commun 5, 4029 (2014).

27. Guler JL, White J, 3rd, Phillips MA, Rathod PK. Atovaquone tolerance in Plasmodium falciparum parasites selected for high-level resistance to a dihydroorotate dehydrogenase inhibitor. Antimicrob Agents Chemother 59, 686–689 (2015).

28. Zheng Y, et al. A Broad Antiviral Strategy: Inhibitors of Human DHODH Pave the Way for Host-Targeting Antivirals against Emerging and Re-Emerging Viruses. Viruses 14, (2022).

29. Davis JP, Cain GA, Pitts WJ, Magolda RL, Copeland RA. The immunosuppressive metabolite of leflunomide is a potent inhibitor of human dihydroorotate dehydrogenase. Biochemistry 35, 1270–1273 (1996).

30. Weisberg AS, et al. Enigmatic origin of the poxvirus membrane from the endoplasmic reticulum shown by 3D imaging of vaccinia virus assembly mutants. Proc Natl Acad Sci U S A 114, E11001–E11009 (2017).

31. Grosenbach DW, et al. Oral Tecovirimat for the Treatment of Smallpox. N Engl J Med 379, 44–53 (2018).

32. Cundy KC, et al. Clinical pharmacokinetics of cidofovir in human immunodeficiency virus-infected patients. Antimicrob Agents Chemother 39, 1247–1252 (1995).

33. GlaxoSmithKline. Product Monograph, PrMEPRON, Atovaquone Oral Suspension, USP 750 mg/5 mL. https://cagskcom/media/6194/mepronpdf, (2016).

34. Gutman J, et al. Mefloquine pharmacokinetics and mefloquine-artesunate effectiveness in Peruvian patients with uncomplicated Plasmodium falciparum malaria. Malar J 8, 58 (2009).

35. Painter WP, et al. Human Safety, Tolerability, and Pharmacokinetics of Molnupiravir, a Novel Broad-Spectrum Oral Antiviral Agent with Activity Against SARS-CoV-2. Antimicrob Agents Chemother, (2021).

36. Kim KS, et al. A quantitative model used to compare within-host SARS-CoV-2, MERS-CoV, and SARS-CoV dynamics provides insights into the pathogenesis and treatment of SARS-CoV-2. PLoS Biol 19, e3001128 (2021).

37. Pittman PRM, J. W.; Kingebeni, P. M.; Tamfum, J-J. M.;Wan, Q.; Reynolds, M. G.; Quinn, X.; Norris, S.; Townsend, M. B.; Satheshkumar, P. S.; Soltis, B.; Honko, A.; Güereña, F. B.; Korman, L.; Huggins, J. W.; The Kole Human Monkeypox Infection Study Group. Clinical characterization of human monkeypox infections in the Democratic Republic of the Congo. medRvix, https://www.medrxiv.org/content/10.1101/2022.1105.1126.22273379v22273371 (2022).

38. Koizumi Y, et al. Quantifying antiviral activity optimizes drug combinations against hepatitis C virus infection. Proc Natl Acad Sci U S A 114, 1922–1927 (2017).

39. Shen L, et al. Dose-response curve slope sets class-specific limits on inhibitory potential of anti-HIV drugs. Nat Med 14, 762–766 (2008).

40. Greco WR, Bravo G, Parsons JC. The search for synergy: a critical review from a response surface perspective. Pharmacol Rev 47, 331–385 (1995).

41. Koizumi Y, Iwami S. Mathematical modeling of multi-drugs therapy: a challenge for determining the optimal combinations of antiviral drugs. Theor Biol Med Model 11, 41 (2014).

42. Jacquerioz FA, Croft AM. Drugs for preventing malaria in travellers. Cochrane Database Syst Rev, CD006491 (2009).

43. Schnellrath LC, Damaso CR. Potent antiviral activity of brequinar against the emerging Cantagalo virus in cell culture. Int J Antimicrob Agents 38, 435–441 (2011).

44. Chan V, Charles BG, Tett SE. Population pharmacokinetics and association between A77 1726 plasma concentrations and disease activity measures following administration of leflunomide to people with rheumatoid arthritis. Br J Clin Pharmacol 60, 257–264 (2005).

45. Johnson RF, et al. Small particle aerosol inoculation of cowpox Brighton Red in rhesus monkeys results in a severe respiratory disease. Virology 481, 124–135 (2015).

46. Saijo M, et al. LC16m8, a highly attenuated vaccinia virus vaccine lacking expression of the membrane protein B5R, protects monkeys from monkeypox. J Virol 80, 5179–5188 (2006).

47. Ohashi H, et al. Identification of Anti-Severe Acute Respiratory Syndrome-Related Coronavirus 2 (SARS-CoV-2) Oxysterol Derivatives In Vitro. Int J Mol Sci 22, (2021).

