## Supplementary information for "Potential anti-monkeypox virus activity of atovaquone, mefloquine, and molnupiravir, and their potential use as treatments"

**Table of contents**

**Supplementary Note-1, 2**
**Supplementary References**
**Fig. S1, S2, S3, S4**
**Table S1, S2, S3, S4**

### Supplementary Note-1:

#### Selection of drugs from the screening

The screening by MPXV infection assay identified 21 drugs that rescued cells from MPXV-induced cytopathology more than 20 fold compared with DMSO-treated control: posaconazole, clotrimazole, didanosine, butoconazole, bifonazole, pyrimethamine, fenticonazole, tolnaftate, isoconazole, miconazole, atovaquone, butenafine, sertaconazole, sulconazole, nelfinavir, mefloquine, amodiaquine, maduramycin, imidocarb, dasabuvir, and molnupiravir.

We selected the focused drugs in the following criteria: We exclude the drugs either 1) that are usually used as topical medications (clotrimazole, butoconazole, bifonazole, fenticonazole, tolnaftate, isoconazole, miconazole, butenafine, sertaconazole, and sulconazole), 2) that are veterinary medicine (maduramycin and imidocarb), or 3) that have been discontinued in manufacture (didanosine, nelfinavir, and dasabuvir). Among the remaining 6 candidates (posaconazole, pyrimethamine, atovaquone, mefloquine, amodiaquine, and molnupiravir), we selected drugs 4) reportedly achieving more than 1 μM of $C_{max}$ in patients as those having preferable pharmacokinetics (pyrimethamine, atovaquone, mefloquine, and molnupiravir). We then selected drugs that reduced intracellular MPXV DNA to less than 50% upon treatment at 3.3 μM for 24 h in the infection assay (atovaquone, mefloquine, molnupiravir) (Fig. S1B) and focused on these three drugs in this study.

### Supplementary Note-2:

#### Quantifying dose-response curve of single drugs

The typical dose-response curves of a single antiviral drug can be analyzed using the following Hill function [18,41,48]:

$$f_u = \frac{1}{1+\left(\frac{D}{IC_{50}}\right)^m}. \quad (1)$$

Here, $f_u$ represents the fraction of infection events unaffected by the drug (i.e., $1-f_u$ equals the fraction of drug-affected events). $D$ is the drug concentration, $IC_{50}$ is the drug concentration that achieves 50% inhibition of activity (i.e., 50% inhibitory concentration), and $m$ is the slope of the dose-response curve (i.e., Hill coefficient). Least-square regression approach was used to fit Eq.(1) to dose-response data and estimate the values of $IC_{50}$ and $m$ as well as $IC_{90}$ (90% inhibitory concentration). Those estimated values for each drug against MPXV are summarized in Table S2.

#### Synergy analysis of drug combination

To evaluate the effect of double-drug combinations, we used a synergy model of Bliss independence [49]. Bliss independence assumes that each drug acts on different targets/mechanisms independently, and the expected combination response is derived from the probabilistic independence of the monotherapy effects as follows:

$$Y_{Bliss} = 1 - (1 - Y_A)(1 - Y_B), \qquad (2)$$

where $Y_A$ and $Y_B$ are the responses of drugs A (i.e., atovaquone) and B (i.e., tecovirimat) that ranges from 0 to 1. Note that a higher value represents a better efficacy of drug. Using Eq. (2), we can calculate the multi-drug synergy score ($S$) for the observed combination response ($Y_{Com}$) as follows:

$$S = Y_{Com} - Y_{Bliss} = Y_{Com} - \big(1 - (1 - Y_A)(1 - Y_B)\big).$$

The synergy scores for the anti-MPXV effects of combined drugs A and B are described in Fig. 6C.

#### Mathematical modeling for the impact of antivirals on MPVX infection in clinical settings

Based on a standard viral dynamics (VD) model, to describe MPXV dissemination among susceptible target cells, we used the following simple mathematical model proposed in the previous papers [18,36,50]:

$$\frac{df(t)}{dt} = -\beta f(t)V(t), \qquad (3)$$

$$\frac{dV(t)}{dt} = \gamma f(t)V(t) - \delta V(t), \qquad (4)$$

where $f(t)$ and $V(t)$ are the ratio of uninfected target cells and the amount of virus, respectively. The parameters $\beta$, $\gamma$, and $\delta$ represent the rate constant for virus infection, the maximum rate constant for viral replication and the death rate of infected cells, respectively.

To quantitatively assess MPXV infection dynamics, we performed a nonlinear least-square fit for the previously reported mean values of the viral load (genomes/ml) in the blood [37] using the following objective function:

$$SSR = \sum_{i=1}^{22} \{\log V(t_i) - \log V^D(t_i)\}^2$$

where $V(t_i)$ and $V^D(t_i)$ are the model-predicted values for the viral load and measured viral load, respectively. The viral load dynamics that were produced with the best fit parameter values are shown in Fig. S3, and the estimated parameters and initial values used here are summarized in Table S4. In the main text, we used the estimated parameters for quantitative simulations. In addition, if we set the threshold of viral load at infection is 0.01 (genomes/ml), the time of the infection, $t_s$, was identified by means of back-calculation when the viral load reaches the threshold. We note that estimated $-t_s = 5.57$ corresponds to the incubation period and found that the period of the best-fit solution is consistent with the published estimates (mean: 8.5 days, 95% credible intervals: 6.6-10.9 days) [51], meaning a validation of our estimation.

To investigate the expected outcome for anti-MPXV treatment with single drugs, we conducted *in silico* experiments with the following mathematical model for post-entry phases such as tecovirimat, cidofovir, atovaquone and molnupiravir;

$$\frac{df(t)}{dt} = -\beta f(t)V(t), \quad (7)$$

$$\frac{dV(t)}{dt} = \big(1 - \varepsilon(t) \times H(t)\big)\gamma f(t)V(t) - \delta V(t), \quad (8)$$

and for entry inhibitor such as mefloquine;

$$\frac{df(t)}{dt} = -\big(1 - \eta(t) \times H(t)\big)\beta f(t)V(t), \quad (9)$$

$$\frac{dV(t)}{dt} = \big(1 - \eta(t) \times H(t)\big)\gamma f(t)V(t) - \delta V(t). \quad (10)$$

Here $H(t)$ is a Heaviside step function defined as $H(t) = 0$ if $t < t^*$: otherwise $H(t) = 1$, where $t^*$ is the initiation of the treatment, and the anti-MPXV effect for $t > t^*$ are described as

$$\varepsilon(t)\ \big(\text{or}\ \eta(t)\big) = 1 - f_u\big(D(t)\big) = 1 - \frac{1}{1 + \left(\frac{D(t)}{IC_{50}}\right)^m},$$

$$D(t) = C_{max} e^{-kt} \quad (t_i < t < t_{i+1}), \quad (11)$$

where $C_{max}$ and $k$ are the maximum drug concentration and the degradation rate for corresponding drug, respectively. $D(t)$ is the concentration of drugs in plasma between $i$-th and $(i+1)$-th doses for $i = 1, \dots, N$. Here $N$ is the number of drug administration, $t_1 = t^*$ is the initiation of the treatment, $t_{i+1} - t_i = \tau$ is the dosing interval, and $t_{N+1} = \infty$ (see Table S3).

#### Evaluation of outcomes for anti-MPXV treatments

To evaluated the outcomes for anti-MPXV treatments, we calculated "cumulative viral load" and "duration of viral shedding" by Eqs. (7-11) (see Fig. S2). The cumulative viral load, i.e., the area under the curve of viral load is defined by $\int_{t_s}^{T_D} V(s)ds$ where $T_D$ is the time for MPXV achieved the detection limit. The duration of viral shedding is calculated as $T_D - t_s$. The detection limit was set to 4.07 genomes/ml which was the smallest value of the viral load in the blood.

### Supplementary References

48. Ohashi H, *et al.* Different efficacies of neutralizing antibodies and antiviral drugs on SARS-CoV-2 Omicron subvariants, BA.1 and BA.2. *Antiviral Res* **205**, 105372 (2022).
49. Zheng S, *et al.* SynergyFinder Plus: Toward Better Interpretation and Annotation of Drug Combination Screening Datasets. *Genomics Proteomics Bioinformatics*, (2022).
50. Iwanami S, *et al.* Detection of significant antiviral drug effects on COVID-19 with reasonable sample sizes in randomized controlled trials: A modeling study. *PLoS Med* **18**, e1003660 (2021).
51. Miura F, *et al.* Estimated incubation period for monkeypox cases confirmed in the Netherlands, May 2022. *Euro Surveill* **27**, (2022).

Fig. S1

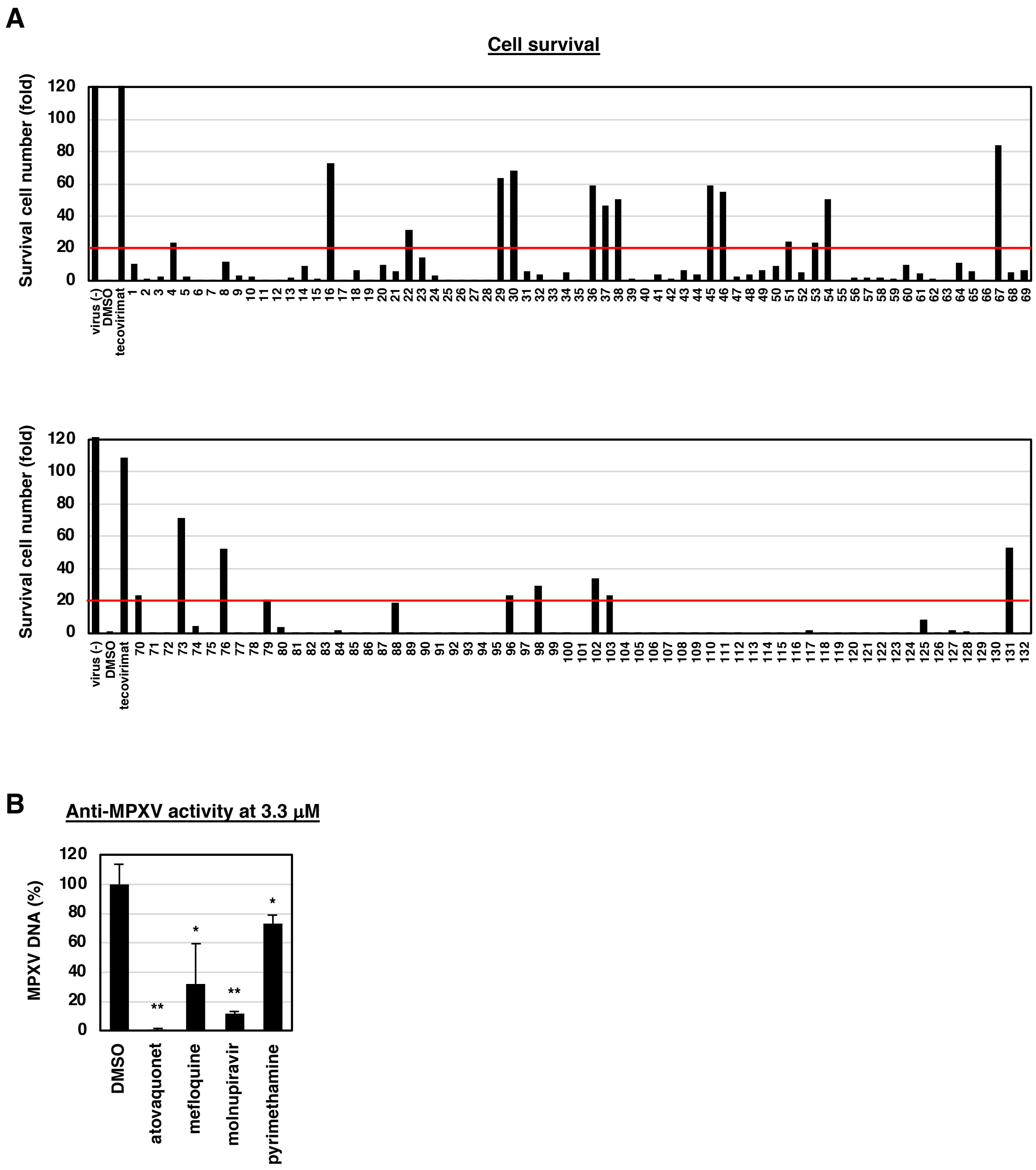


**Fig. S1. Screening of drug library in MPXV-infected VeroE6 cells.** (A) VeroE6 cells treated as shown in Fig. 1A were fixed and stained with DAPI to quantify survival cell number by a high-content imaging analyzer. MPXV infection upon DMSO treatment reduced cell survival to less than 1%, compared with uninfected cells. The red line shows the 20-fold cell survival reltive to DMSO-control in MPXV-infected cells. The names of the drugs for each number are described in Table S1. (B) As the second screening, VeroE6 cells infected with MPXV were treated with the drugs at 3.3 μM for 24 h to quantify intracellular MPXV DNA by real time PCR.

Fig. S2

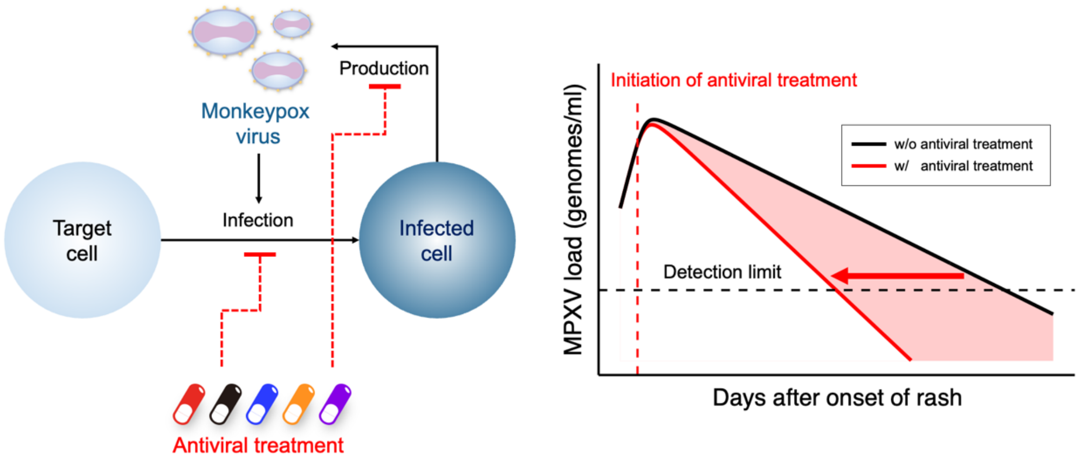


**Fig. S2. Schematics of the model for predicting the impact of drugs on virus infection dynamics in patients.** Typical MPXV load in patients undergoing antiviral treatment is shown. The outcomes for treatment, that is, reduction in “cumulative viral load” (area under the curve for MPXV load, right graph) and “duration of viral shedding” (red arrow, right graph) are graphically depicted.

Fig. S3

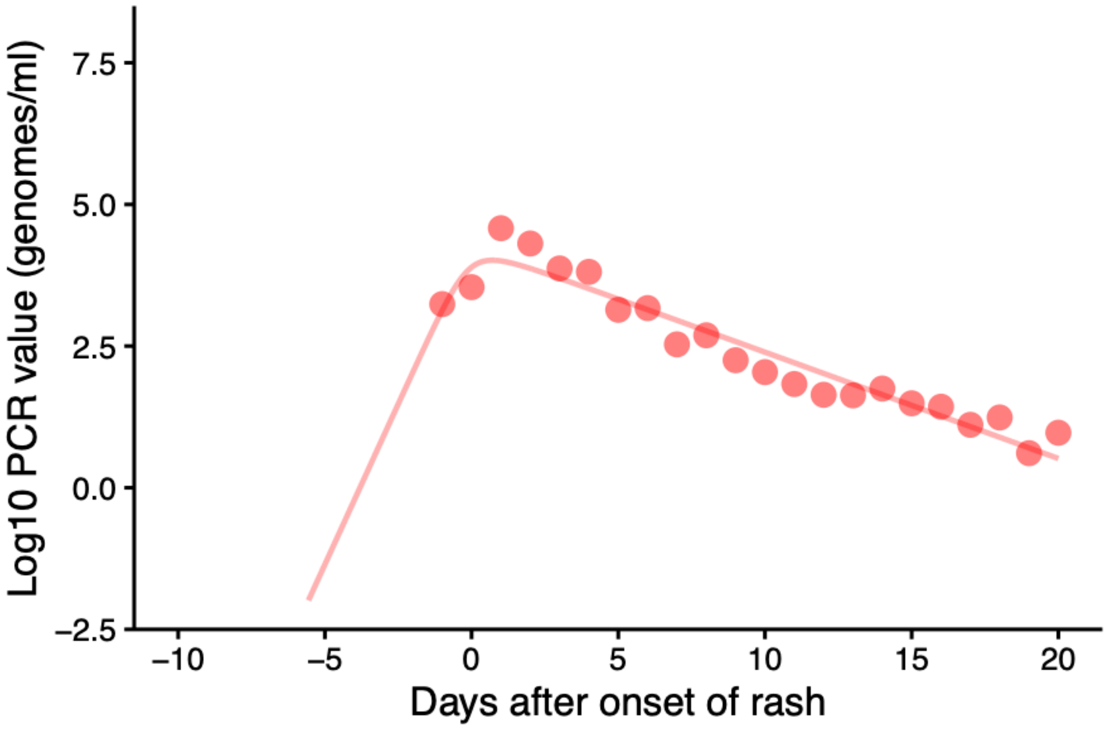


**Fig. S3. Expected MPXV infection dynamics.** The predicted variability of the MPXV load dynamics is shown based on a nonlinear least-square estimation for the previously reported mean values of the viral load (genomes/ml) [37]. The solid line gives the best-fit solution for Eqs.(3-4), and the dots show the mean values of MPXV loads from blood.

Fig. S4

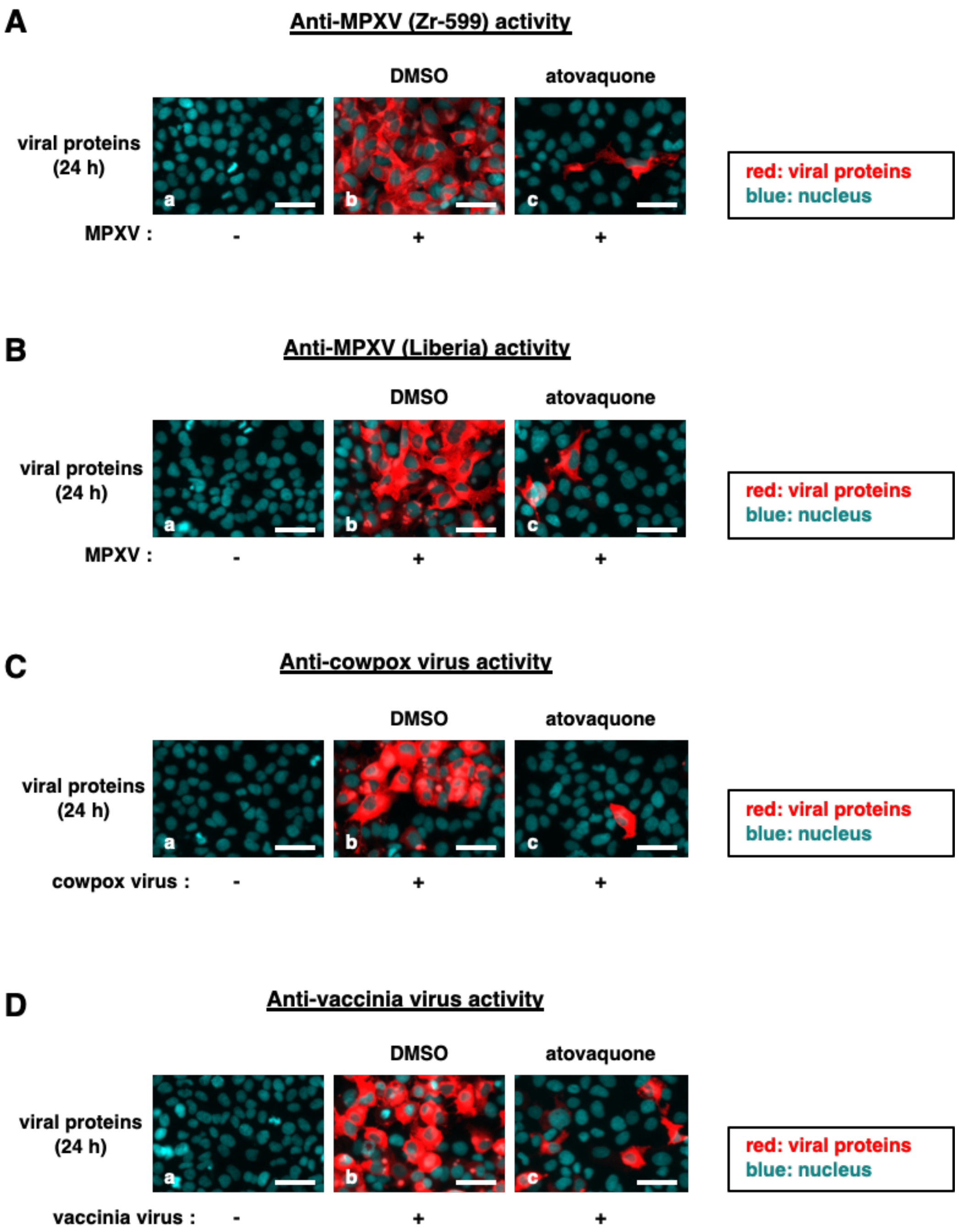


**Fig. S4. Anti-MPXV activity of atovaquone in Huh-7 cells.** A human hepatoma-derived Huh-7 cells were inoculated with MPXV (Zr-599, Liberia strains), vaccinia virus, or cowpox virus at the same amount of the inoculum and with the same treatment time as used in Fig. 7. Viral proteins (red) as well as the nuclei (blue) were detected as shown in Fig. 7.

**Table. S1. List of drugs in the library.**

| | |
|---|---|
| Danoprevir | 1 |
| Ritonavir | 2 |
| Entecavir Hydrate | 3 |
| Posaconazole | 4 |
| Artemisinin | 5 |
| Fluconazole | 6 |
| Ivermectin | 7 |
| Lopinavir | 8 |
| Stavudine | 9 |
| Tenofovir Disoproxil Fumarate | 10 |
| Voriconazole | 11 |
| Atazanavir Sulfate | 12 |
| Daclatasvir | 13 |
| Natamycin | 14 |
| Telaprevir | 15 |
| Clotrimazole | 16 |
| Darunavir Ethanolate | 17 |
| Amprenavir | 18 |
| Telbivudine | 19 |
| Flucytosine | 20 |
| Amorolfine HCl | 21 |
| Didanosine | 22 |
| Emtricitabine | 23 |
| Lamivudine | 24 |
| Zalcitabine | 25 |
| Terbinafine | 26 |
| Nevirapine | 27 |
| Aciclovir | 28 |
| Butoconazole nitrate | 29 |
| Bifonazole | 30 |
| Valaciclovir HCl | 31 |

| | |
|---|---|
| Ganciclovir | 32 |
| Rimantadine | 33 |
| Elvitegravir | 34 |
| Raltegravir | 35 |
| Pyrimethamine | 36 |
| Fenticonazole Nitrate | 37 |
| Tolnaftate | 38 |
| Artemether | 39 |
| Amantadine HCl | 40 |
| Famciclovir | 41 |
| Pyrantel Pamoate | 42 |
| Quinine HCl Dihydrate | 43 |
| Ribavirin | 44 |
| Isoconazole nitrate | 45 |
| Miconazole | 46 |
| Zidovudine | 47 |
| Dolutegravir | 48 |
| Sofosbuvir | 49 |
| Clevudine | 50 |
| Atovaquone | 51 |
| Ornidazole | 52 |
| Butenafine HCl | 53 |
| Sertaconazole nitrate | 54 |
| Moxidectin | 55 |
| Velpatasvir | 56 |
| Grazoprevir | 57 |
| Boceprevir | 58 |
| Benznidazole | 59 |
| Lumefantrine | 60 |
| Quinidine sulfate | 61 |
| Arteether | 62 |

| | |
|---|---|
| Valganciclovir HCl | 63 |
| Ronidazole | 64 |
| Tinidazole | 65 |
| Zinc Pyrithione | 66 |
| Sulconazole Nitrate | 67 |
| Climbazole | 68 |
| Penciclovir | 69 |
| Broxyquinoline | 70 |
| Primaquine Diphosphate | 71 |
| Luliconazole | 72 |
| Nelfinavir Mesylate | 73 |
| Anidulafungin | 74 |
| Bephenium Hydroxynaphthoate | 75 |
| Mefloquine HCl | 76 |
| Ethylparaben | 77 |
| Diethylcarbamazine citrate | 78 |
| Amodiaquine dihydrochloride dihydrate | 79 |
| Clioquinol | 80 |
| Atazanavir | 81 |
| Efavirenz | 82 |
| Nicarbazin | 83 |
| Neticonazole Hydrochloride | 84 |
| Asunaprevir | 85 |
| Tavaborole | 86 |
| Brivudine | 87 |
| Simeprevir | 88 |
| Isoprinosine | 89 |
| Efinaconazole | 90 |
| Terconazole | 91 |
| Milbemycin Oxime | 92 |
| Daclatasvir Digydrochloride | 93 |
| Ganciclovir sodium | 94 |

| | |
|---|---|
| Febantel | 95 |
| Maduramycin Ammonium | 96 |
| Doramectin | 97 |
| Imidocarb dipropionate | 98 |
| Abacavir | 99 |
| Raltegravir potassium | 100 |
| Selamectin | 101 |
| Tizoxanide | 102 |
| Dasabuvir | 103 |
| Ombitasvir | 104 |
| Paritaprevir | 105 |
| Quinacrine Dihydrochloride Dihydrate | 106 |
| Dimetridazole | 107 |
| Amenamevir | 108 |
| Elbasvir | 109 |
| Myclobutanil | 110 |
| Glecaprevir | 111 |
| Ledipasvir | 112 |
| Cabotegravir | 113 |
| Tenofovir Alafenamide | 114 |
| Favipiravir | 115 |
| Indinavir Sulfate | 116 |
| Cidofovir (2 μM) | 117 |
| Zanamivir (2 μM) | 118 |
| Chloroquine Phosphate (2 μM) | 119 |
| Hydroxychloroquine Sulfate (2 μM) | 120 |
| Foscarnet Sodium (2 μM) | 121 |
| Caspofungin Acetate (2 μM) | 122 |
| Micafungin Sodium (2 μM) | 123 |
| Ketoconazole | 124 |
| Itraconazole | 125 |

| | |
|---|---|
| Oseltamivir Phosphate | 126 |
| Flubendazole | 127 |
| Clopidol | 128 |
| Piperaquine phosphate (2 μM) | 129 |
| Remdesivir | 130 |
| Molnupiravir | 131 |
| Nirmatrelvir | 132 |

Drugs without concentration were treated at 10 μM, whereas those treated at 2 μM are indicated.

**Table S2.** Estimated characteristic parameters of the tested antiviral drugs

| Drug (unit) | Class | $IC_{50}$ | $IC_{90}$ | $m$ |
|---|---|---|---|---|
| Single drug treatment | | | | |
| Tecovirimat (nM) | PEI | 2.80 | 10.3 | 1.69 |
| Cidofovir (μM) | PEI | 13.7 | 43.8 | 1.90 |
| Atovaquone (μM) | PEI | 0.516 | 0.763 | 5.61 |
| Mefloquine (μM) | EI | 5.19 | 7.85 | 5.32 |
| Molnupiravir (μM) | PEI | 1.35 | 3.25 | 2.49 |

PEI, post entry inhibitor; EI, entry inhibitor
$IC_{50}$, 50% inhibitory concentration
$IC_{90}$, 90% inhibitory concentration
m, slope of the dose-response curve (i.e., Hill coefficient)

**Table S3.** Summary of pharmacokinetic parameters of anti-MPXV drugs.

| Parameter name | Symbol | Unit | Tecovirima | Cidofovir | Atovaquone | Mefloquine | Molnupiravir |
|---|---|---|---|---|---|---|---|
| Single compartment model | | | | | | | |
| Maximum concentration | $C_{max}$ | μg/ml | 2.21 | 20.7 | 11.5 | 2.700 | 3.64 |
| Degradation rate | $k$ | day$^{-1}$ | 0.723 | 7.56 | 0.248 | 0.0479 | 12.9 |
| Molecular mass | – | g/mol | 394.35 | 315.22 | 366.84 | 414.78 | 259.22 |
| Dosing schedule | | | | | | | |
| Initiation of treatment | $t^*$ | days after onset of rash | -1 | -1 | -1 | -1 | -1 |
| Dosing interval | $\tau$ | day | 0.5 | 7 | 0.5 | --- | 0.5 |
| Number of drug administration | $N$ | --- | 28 | 2 | 42 | 1 | 10 |

Tecovirimat: 600 mg, PO BID for 14 days [31]
Cidofovir: 5 mg/kg, IV qWeek × 2 doses [32]
Atovaquone: 750 mg, PO BID for 21 days [33]
Mefloquine: 25 mg/kg, PO once with artesunate [34]
Molnupiravir: 800 mg, PO BID for 5 days [35]

**Table S4.** Estimated parameters and initial values for MPXV infection

| Parameter name | Symbol | Unit | Value |
|---|---|---|---|
| Maximum rate constant for viral replication | $\gamma$ | day$^{-1}$ | 1.37 |
| Rate constant for virus infection | $\beta$ | (copies/ml)$^{-1}$ day$^{-1}$ | 1.70×10$^{-4}$ |
| Death rate of infected cells | $\delta$ | day$^{-1}$ | 0.435 |
| Initial viral load | $V(0)$ | genomes/ml | 7.78×10$^{3}$ |
| Calculated values | | | |
| Time of infection | $t_s$ | day | -5.57 |
| Fraction of target cells at time of infection | $f(t_s)$ | --- | 2.22 |
| Viral load at time of infection* | $V(t_s)$ | genomes/ml | 0.01 |

* Threshold of viral load at infection is fixed to be 0.01
